# Changes in freshwater macroinvertebrate richness due to river impoundment in the United States

**DOI:** 10.1101/814335

**Authors:** Gabrielle Trottier, Katrine Turgeon, Francesca Verones, Daniel Boisclair, Cécile Bulle, Manuele Margni

**Affiliations:** CIRAIG, Polytechnique Montréal, Département de mathématiques et génie industriel, C.P. 6079, succ. Centre-Ville, Montréal, Québec, Canada, H3C 3A7; ISFORT, Université du Québec en Outaouais, 58 rue Principale, Ripon, Québec, Canada, J0V1V0; Industrial Ecology Program, Department of Energy and Process Engineering, NTNU, Sem Sælands vei 7, Trondheim, Norway, 7491; Département des sciences biologiques, Université de Montréal, C.P. 6128 Succursale « Centre-ville », Montréal, Québec, H3C 3J7, Canada; Département de stratégie, responsabilité sociale et environnementale, École des sciences de la gestion, Université du Québec à Montréal, Montréal, Québec, Canada, H3C 3P8

**Keywords:** Life cycle assessment, Reservoirs, Biodiversity, Macroinvertebrates, Water management, Aquatic ecology

## Abstract

Whether it is for water supply, flood control or hydropower uses, the transformation of a river into a reservoir can impact freshwater ecosystems and their biodiversity. Using the National Lake Assessment (NLA; 148 reservoirs) and the National Rivers and Streams Assessment (NRSA; 2121 rivers and streams) of the United States Environmental Protection Agency (USEPA), we evaluated the impacts of river impoundment on macroinvertebrate biodiversity at three spatial scales (*i.e.*, reservoir, ecoregion and country scale). We used a space-for-time substitution approach to model the impact of impoundment (*i.e.*, we used rivers and streams as the before-impoundment conditions, and reservoirs as the after-impoundment conditions). We expressed the impact on biodiversity in terms of potentially disappeared fraction of species (PDF) to be used in the life cycle assessment (LCA) framework. To understand the role of regionalization, and some potentially influential variables, on changes in macroinvertebrate richness following impoundment in the United States, we used analyses of variance (ANOVAs) as well as variation partitioning, and developed empirical predictive models. Overall, 26% of macroinvertebrate taxa disappeared following impoundment in the United States, and PDFs followed a longitudinal gradient across ecoregions (*i.e.*, higher PDFs in the western part of the country, lower PDFs in the eastern part). We also observed that large and oligotrophic reservoirs, located in high elevation had high PDFs. This study provides the first empirical PDF values for macroinvertebrates to be used as characterization factors (CFs) by LCA practitioners. We also provide strong support for regionalization and a simple predictive model to be used by LCA modellers.

## 1. INTRODUCTION

Population growth in the last century has inevitably resulted in an increased need, and use of water by humans (WCD 2000, Strayer and Dudgeon 2010, Fang and Jawitz 2019). The extraction of water and the regulation of water flow by dams (*i.e.*, creation of storage reservoirs for drinking water, flood control, and energy production) or the water diversion by channels (*i.e.*, irrigation and navigation) has benefited human populations across the globe (Richter et al. 2003, Lehner et al. 2011, Chen et al. 2016). Despite some clear societal benefits, water use is often accompanied by a myriad of environmental impacts on freshwater ecosystems (Rosenberg et al. 2000, Vörösmarty 2005, Abell et al. 2008, Renöfält et al. 2010, Gracey and Verones 2016).

The environmental impacts of a dam, and of transforming a river into a reservoir are particularly well documented (Agostinho et al. 2008). The geomorphology of a reservoir is different from the one of the original river. The water depth, as well as the hydrological regime (*i.e.*, lentic instead of lotic system), are notably altered. Water depth is known to affect ecological community structure and productivity, alongside with total dissolved solids (Ryder 1965, 1982, Ryder et al. 1974, Youngs and Heimbuch 1982, Jackson et al. 1990, Rempel and Colby 1991). A change in the hydrological regime affects several physical and biological processes (erosion/sedimentation; biological cue to life-history strategies), and the organisms’ capacity to thrive and survive in these ecosystems. Those changes can ultimately impact ecosystem biodiversity, productivity, and the provision of ecosystem services (Poff et al., 1997; Bunn & Arthington, 2002; Dudgeon et al., 2006). If we wish to sustain these services and preserve the ecological integrity of freshwater ecosystems, it is imperative to understand the potential impacts of dams and river impoundment on freshwater biodiversity.

The life cycle assessment (LCA) approach allows the evaluation of the potential environmental impacts of a product, process, or service throughout its entire life cycle (*i.e.*, from resource extraction to end of life; ISO, 2006). Emissions and resource extraction (*i.e.*, inventory flows) related to all activities involved in the life cycle of a product, process, or service are first inventoried and then characterized into potential environmental impacts through so-called characterization factors (CFs) (Curran et al. 2011). CFs are then modeled up to damages on different areas of protection, from an anthropocentric point of view, and traditionally include ecosystem quality, human health, and natural resources (Verones et al. 2017). The ecosystem quality area of protection typically aggregates multiple damage level indicators, such as eutrophication, acidification, ecotoxicity, as well as damages from climate change, water use and land use. Damage level indicators regarding the ecosystem quality area of protection are traditionally expressed in PDF·m^2^·yr units, where PDF stands for potentially disappeared fraction of species, and can be aggregated into one common damage level indicator or kept disaggregated (Verones et al., 2017, Milà i Canals et al., 2007).

The impacts of transforming a river into a reservoir on ecosystem quality have received little attention in LCA. To our knowledge, only few attempts to evaluate changes in fish richness (*i.e.*, the number of taxa present) following impoundment have been made (see Turgeon et al., 2019 [*submitted*], available as a preprint and Dorber et al., 2019). In this study, we assessed the potential impacts of impoundment (*i.e.*, transforming a river into a reservoir) using empirical data on changes in freshwater macroinvertebrate richness based on a dataset of 148 reservoirs and 2121 rivers and streams across the continental United States. Using a space-for-time substitution, we derived PDFs at three spatial scales: the scale of the United States, the scale of nine ecoregions, and the scale of reservoirs. Finally, we developed an empirical predictive model, built from the most influential variables affecting PDFs at the reservoir scale, to be used by LCA modellers.

## 2. MATERIAL AND METHODS

### 2.1 Impact characterization framework

CFs are determined by characterization models. These models are either based on the modelling of the environmental mechanisms that link environmental interventions to impact indicators, or on empirical observations of environmental interventions on impact indicators. Two categories of damage level indicators exist for the ecosystem quality area of protection. Traditionally, ecosystem quality damage assessment models used to quantify the temporary disappearance of species integrated in time and space, whether this was expressed in PDF·m^2^·yr (*i.e.*, a potentially disappeared fraction of species over a given area and duration; Jolliet et al., 2003) or in species·yr (Goedkoop et al. 2009).

More recently, some models focused on the regional or global disappearance of species expressed in PDF, that is quantifying the permanent disappearance of species at the continental or global scale (de Baan et al. 2013, Chaudhary et al. 2015). Both categories of indicators are relevant and should be considered complementary. The first one allows to assessment of temporary degradation of an ecosystem that will ultimately recover (*e.g.*, a toxic contamination that ultimately is degraded and after which the affected ecosystem recovers). The second one allows the assessment of the absolute disappearance of species that will never be recovered (*e.g.*, endemic species that disappeared because of land transformation). The indicator developed in the current research corresponds to the first category of indicators and is meant to quantify the temporary damage on freshwater ecosystems, more specifically macroinvertebrates, due to impoundment, and integrated in time and in space.

The framework proposed by de Baan (2013) and Chaudhary (2015) for land occupation has been adopted and adapted to assess the occupation of a water body. In this approach, the indicator builds on an empirical model assessing land use impacts on biodiversity, and is expressed in PDF·m^2^·yr. In this case, the CF is expressed in PDF units, or implicitly PDF·m^2^·yr/m^2^·yr of water body occupied. This CF is the observed change in taxa richness, with respect to a reference ecosystem, and multiplies the environmental intervention, which is expressed in terms of m^2^·yr of water body occupied. In the current research, we did not quantify the damage due to water body transformation but only the damage of water body occupation. To be able to quantify the damage of water body transformation in compliance with the Milà i Canals framework (2007), the time needed by the water body to recover after the impoundment ends would be needed and we don’t have this information. That’s why we limited ourselves to the assessment of the impacts of occupation.

Here, we used an empirical space-for-time substitution approach to examine the difference in macroinvertebrate richness between a natural reference ecosystem and an impacted system to generate empirical PDFs representing the impacts of occupying a river, which has been transformed into a reservoir. To calculate CFs, we used the PDF; or implicitly PDF·m^2^·yr/m^2^·yr of water body occupied) as a unit to express the proportional loss of macroinvertebrate taxa following impoundment, specific to nine ecoregions and 148 reservoirs in the United States.

### 2.2 Data collection

#### 2.2.1 Macroinvertebrate richness

To extract data on macroinvertebrate richness in reservoirs (impacted sites; after impoundment), we used the 2012 National Lake Assessment (NLA), which is a United States Environmental Protection Agency (USEPA) effort to statistically survey numerous ponds, lakes and reservoirs in the United States and their associated biological, chemical, physical and recreational characteristics (USEPA 2015a). From this dataset, we retrieved macroinvertebrates richness (*i.e.*, RICHNESS; taxonomic resolution at the generic level, except for oligochaetes, mites, and polychaetes, which were identified to the family level, and ceratopogonids at the subfamily level; USEPA, 2017), a unique identifier (UID) for each reservoir, the latitude (LAT), longitude (LON), the ecoregion (ECO), and a suite of environmental variables (Table 1) from 148 reservoirs across the United States (Figure 1, reservoirs shown in black). It was impossible to get macroinvertebrate richness for the pre-impoundment period for the same ecosystem. To get an estimate of macroinvertebrates taxa richness from pre-impoundment conditions, we used the space-for-time substitution approach (Pickett 1989) and used the 2008-2009 National Rivers and Streams Assessment (NRSA). This dataset is also a USEPA initiative to survey U.S. rivers and streams’ biological, chemical, physical, and recreational characteristics (USEPA 2015b). The NRSA dataset serves as reference for pre-impoundment characteristics, given the assumption that taxa richness in rivers and streams in the surrounding area of a reservoir from the NLA dataset would be comparable to what would have been found in a river prior to its transformation into a reservoir. The same variables were collected (UID, LAT, LON, ECO and RICHNESS) for 2121 rivers and streams across the United States (Figure 1, rivers and streams shown in white).

**Figure 1.**
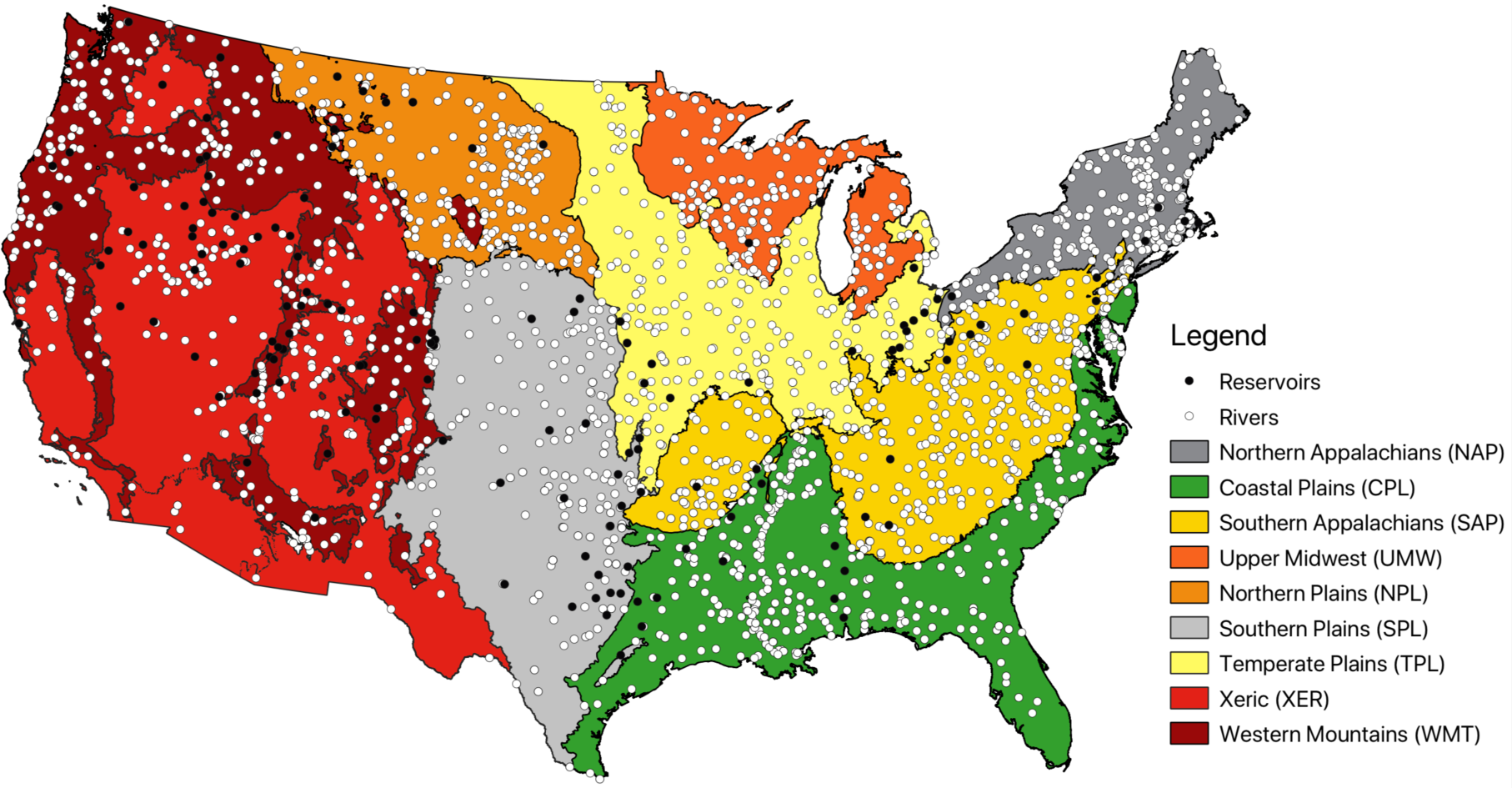
Map of the USEPA-NLA, where reservoirs (*n* = 148) and the USEPA-NRSA rivers and streams data points (*n* = 2121), color-coded according to ecoregions.

**Table 1.** Table showing the four matrices used for variation partitioning, as well as the variables they included, a short definition, their respective units and the type of variable they are (*i.e.*, N for numerical and F for categorical). Variables in bold are the variables selected for the parsimonious variation partitioning.

| MATRIX | VARIABLE | DEFINITION | UNITS | TYPE |
| --- | --- | --- | --- | --- |
| SPATIAL | LATITUDE | LATITUDE OF RESERVOIR | DECIMAL DEGREES | N |
|  | LONGITUDE | LONGITUDE OF RESERVOIR | DECIMAL DEGREES | N |
|  | **ECOREGION** | NARS 9-LEVEL REPORTING REGIONS, BASED ON AGGREGATED OMERNIK LEVEL III ECOREGIONS | - | F |
|  | TEMPERATURE | ANNUAL MEAN AIR TEMPERATURE, SPECIFIC TO ECOREGION | °C | N |
|  | PRECIPITATION | ANNUAL MEAN PRECIPITATIONS, SPECIFIC TO ECOREGION | MM | N |
|  | LC.FORESTED | PERCENTAGE OF LAND COVER IN ECOREGION THAT IS FORESTED | % | N |
|  | LC.CULTIVATED.PASTURE | PERCENTAGE OF LAND COVER IN ECOREGION THAT IS CULTIVATED PASTURES | % | N |
|  | LC.WETLANDS | PERCENTAGE OF LAND COVER IN ECOREGION THAT IS WETLANDS | % | N |
|  | LC.GRASSLAND.SHRUBS | PERCENTAGE OF LAND COVER IN ECOREGION THAT IS GRASSLANDS AND SHRUBS | % | N |
|  | LC.DEVELOPED | PERCENTAGE OF LAND COVER IN ECOREGION THAT IS DEVELOPED | % | N |
|  | LC.WATER.BARREN | PERCENTAGE OF LAND COVER IN ECOREGION THAT IS WATER OR BARREN | % | N |
| PHYSICAL | **AREA** | SURFACE AREA OF RESERVOIR | HA | N |
|  | **ELEVATION** | ELEVATION RESERVOIR COORDINATES | M | N |
|  | MACROPHYTES | INDEX OF TOTAL COVER OF AQUATIC MACROPHYTES OF RESERVOIR | - | N |
|  | SHALLOW.WATER | SHALLOW WATER HABITAT CONDITION INDICATOR | - | N |
|  | RIP.VEG | RIPARIAN VEGETATION CONDITION INDICATOR | - | N |

**Table 1. Continued**
| MATRIX | VARIABLE | DEFINITION | UNITS | TYPE |
| --- | --- | --- | --- | --- |
| CHEMICAL | **TROPHIC.STATE** | TROPHIC STATE OF RESERVOIR (*i.e.*, OLIGOTROPHIC AND EUTROPHIC) | - | F |
|  | SECCHI | SECCHI DEPTH | M | N |
|  | DOC | DISSOLVED ORGANIC CARBON LEVEL | MG/L | N |
|  | PTL | TOTAL PHOSPHORUS LEVEL | UG/L | N |
|  | COLOR | WATER COLOR | PCU | N |
|  | COND | WATER CONDUCTIVITY LEVEL | US/CM | N |
|  | NTL | TOTAL NITROGEN LEVEL | MG/L | N |
|  | **PH** | PH LEVEL | PH SCALE | N |
|  | METHYLMERCURY | TOP SEDIMENT METHYLMERCURY LEVEL | NG/L | N |
|  | CHLA | CHLOROPHYLL-A MEASUREMENT RESULT OF RESERVOIR | UG/L | N |
| HUMAN | BUILDINGS | HUMAN INFLUENCE BY BUILDINGS AROUND RESERVOIR SHORELINE | - | N |
|  | COMMERCIAL | HUMAN INFLUENCE BY COMMERCIAL ACTIVITIES AROUND RESERVOIR SHORELINE | - | N |
|  | CROPS | HUMAN INFLUENCE BY CROPS AROUND RESERVOIR SHORELINE | - | N |
|  | DOCKS | HUMAN INFLUENCE BY DOCKS AROUND RESERVOIR SHORELINE | - | N |
|  | LANDFILL | HUMAN INFLUENCE BY LANDFILL AROUND RESERVOIR SHORELINE | - | N |
|  | **LAWN** | HUMAN INFLUENCE BY LAWN AROUND RESERVOIR SHORELINE | - | N |
|  | PARK | HUMAN INFLUENCE BY PARKS AROUND RESERVOIR SHORELINE | - | N |
|  | PASTURE | HUMAN INFLUENCE BY PASTURES AROUND RESERVOIR SHORELINE | - | N |
|  | POWERLINES | HUMAN INFLUENCE BY POWERLINES AROUND RESERVOIR SHORELINE | - | N |
|  | **ROADS** | HUMAN INFLUENCE BY ROADS AROUND RESERVOIR SHORELINE | - | N |
|  | WALLS | HUMAN INFLUENCE BY WALLS AROUND RESERVOIR SHORELINE | - | N |
|  | OTHER | HUMAN INFLUENCE BY OTHER AROUND RESERVOIR SHORELINE | - | N |

#### 2.2.2 Native riverine taxa definition

The taxa pool observed in rivers and streams was used as a baseline (reference; native riverine taxa) to compare taxa richness before and after impoundment. Using only native riverine taxa excluded all new taxa that would be encountered in a lake-like habitat (*i.e.*, reservoir), since they would most likely not be present in a pre-impoundment, river-like habitat. Comparing to a mean reference in each ecoregion, instead of a single river/stream close to the reservoir, ensures that we are measuring the impacts from a set of reference conditions and not a singular pristine, or impacted, river or stream.

#### 2.2.3 Ecoregions

Reservoirs and rivers/streams were distributed across nine terrestrial ecoregions to see if patterns varied across the United States; Coastal Plains (CPL), Northern Appalachians (NAP), Northern Plains (NPL), Southern Appalachians (SAP), Southern Plains (SPL) Temperate Plains (TPL), Upper Midwest (UMW), Western Mountains (WMT) and Xeric (XER; Figure 1). These ecoregions are the result of an aggregation (Omernik, 1987; Herlihy et al., 2008), based on similar environmental characteristics (*i.e.*, climate, vegetation, soil type and geology) and macroinvertebrate assemblages similarity of the level III ecoregions (Omernik 1987), which was adopted by the USEPA for both the NLA and NRSA surveys (USEPA 2016). For each ecoregion, we also extracted some land cover variables (USEPA, 2016; Table 1, spatial matrix) and variables related to human impacts (Table 1, human matrix).

#### 2.2.4 PDFs calculation

We calculated PDFs as the difference in richness between reference river (*x*) and impacted reservoir richness (*y*), divided by the reference river richness (*x*). At the United States scale (*PDF*_*usa*_), we compared overall United States mean native riverine richness in rivers and/or stream (one observation of richness per river and/or stream averaged over the United States; *x̄*_*usa*_) to the overall United States mean richness in reservoirs (one observation of richness per reservoir averaged over the United States; *ӯ*_*usa*_) to get a United States specific change in richness, as per equation 1;

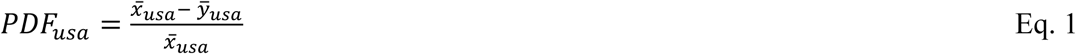

At the ecoregion scale (*PDF*_*eco*_), we compared ecoregion mean native riverine richness of all rivers and/or stream (one observation of richness per river and/or stream averaged over each ecoregion; *x̄*_*eco*_) to the ecoregion mean richness in reservoirs (one observation of richness per reservoir averaged over each ecoregion; *ӯ*_*eco*_) to get an ecoregion specific change in richness, as per equation 2;

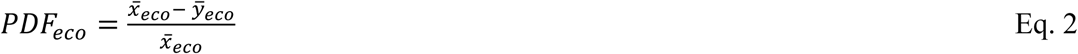

At the reservoir scale (*PDF*_*res*_), we compared ecoregion mean native riverine richness of all rivers and stream (one observation of richness per river and/or stream averaged over each ecoregion; *x̄*_*eco*_) to the richness of a specific reservoir within the same ecoregion (one specific richness observation per reservoir, no averaging; *y*_*res*_) to get a reservoir specific change in richness, as per equation 3;

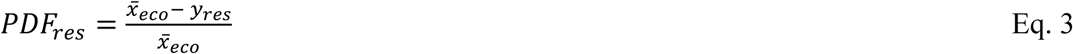

### 2.3 Data analysis and empirical modelling

#### 2.3.1 Regionalization and ANOVA

Regionalization refers to the integration of existing spatial variability to improve results’ representativeness and reduce spatial uncertainties. Regionalization is a critical aspect in LCA (Patouillard et al. 2016, Yang 2016). CFs must be developed at an appropriate scale to capture the environmental impacts of a product, process, or service, and, to inform decision-makers. Here, we examined if PDFs differed across nine terrestrial ecoregions (Herlihy et al. 2008) that were adopted by both the NLA and NRSA surveys. As a first step, we ran an analysis of variance (ANOVA) to determine if PDFs differed across ecoregions (*i.e.*, ecoregion scale) and thus, test the relevance of this regionalization scale. To do so, we used a one-way randomized-group ANOVA. We assessed the significance of regionalization at the ecoregion scale and identified which ecoregions were significantly different from each other based on the standardized mean difference and their confidence intervals (CIs). We conducted the ANOVA with the *ind.oneway.second* function in the *rpsychi* package version 0.8 (Okumura 2012).

#### 2.3.2 Variation partitioning to explain the variation observed in our PDFs

As a second step, we were interested to understand which variables, associated to reservoirs or ecoregions, explained the variation observed in PDFs at the reservoir scale. To do so, we used variation partitioning (Legendre 2008). Variation partitioning describes how a set of explanatory matrices explains the shared variation observed in a response variable (*i.e.*, PDF). We built four explanatory matrices based on the availability of the descriptive variables, and selected relevant variables potentially influencing richness based on expert judgment, from the NLA and NRSA datasets; a spatial matrix (location of the reservoirs*; i.e.,* latitude, ecoregion, and types of land covers), a physical matrix (variables describing the reservoir*; i.e*., reservoir area, elevation), a chemical matrix (biochemical state of the reservoir*; i.e*., pH, trophic state, dissolved organic carbon) and a human activity matrix (human influence around the reservoir shoreline*; i.e*., influence of crops, influence of docks). See Table 1 for a complete description of the variables included in each matrix. To achieve the most parsimonious analysis, we performed a stepwise selection procedure on each explanatory matrix to identify which variable, or combination of variables, best-explained the variation in PDFs (variables in bold; Table 1). Variation partitioning was conducted with the *varpart* function in the *vegan* package version 2.5-2 (Oksanen et al. 2019).

#### 2.3.3 Empirical modelling

As a third step, we developed an empirical predictive model explaining the variation observed in our PDFs to be used by LCA modellers. We combined the variables identified by the variation partitioning analysis as the most influential variables to explain the variation in PDFs from the four above-mentioned explanatory matrices into one matrix and performed a model selection procedure. However, as opposed to the variation partitioning, ecoregion was not used in the empirical model to avoid constraining the model to the ecoregions of the United States and thus, allows extrapolation of the model. We use an automatic forward stepwise model selection procedure, which builds a model by maximizing adjusted R^2^, stopping the selection when it starts to decrease (Oksanen et al. 2019). We used the recommended information theoretic approach based on AIC (Burnham and Anderson 2004) and a Bayesian information criterion (BIC; Schwarz, 1978) to compare the candidate models, and select the model with the highest support. Predictive modelling was performed with the function *ordiR2step*, also from the *vegan* package. All statistical analyses were made using R version 3.0.2 (R Core Team 2013).

## 3. RESULTS

### 3.1 Variation in PDFs across ecoregions and scales

Empirical PDFs showed a loss in macroinvertebrate richness following impoundment in the United States, and this loss followed a longitudinal gradient associated with the ecoregions (Figure 2). At the scale of the United States, 26% of macroinvertebrate taxa disappeared in response to river impoundment (PDF = 0.255 ± 0.123 [mean ± SE]; Table 2 and Figure 2). At the scale of the ecoregion, six out of nine ecoregions showed a significant loss of macroinvertebrate taxa, varying from 0.049 ± 0.048 to 0.472 ± 0.061, and one ecoregion (CPL), showed a significant gain of macroinvertebrate taxa (*i.e.*, -0.242 ± 0.114; Table 2 and Figure 2). PDFs in most ecoregions significantly differed from each other except for NAP, NPL and UMW. Those ecoregions were characterized by small sample sizes (respectively, *n* = 4, 8 or 2; Figure 3). Results from the ANOVA suggest a longitudinal gradient of impact. At the reservoir scale, PDFs varied from -0.995 ± 0.640 (observation ± pooled SD) to 1.000 ± 0.323, and 62% of the reservoirs showed a significant loss of macroinvertebrates taxa (Table A1).

**Figure 2.**
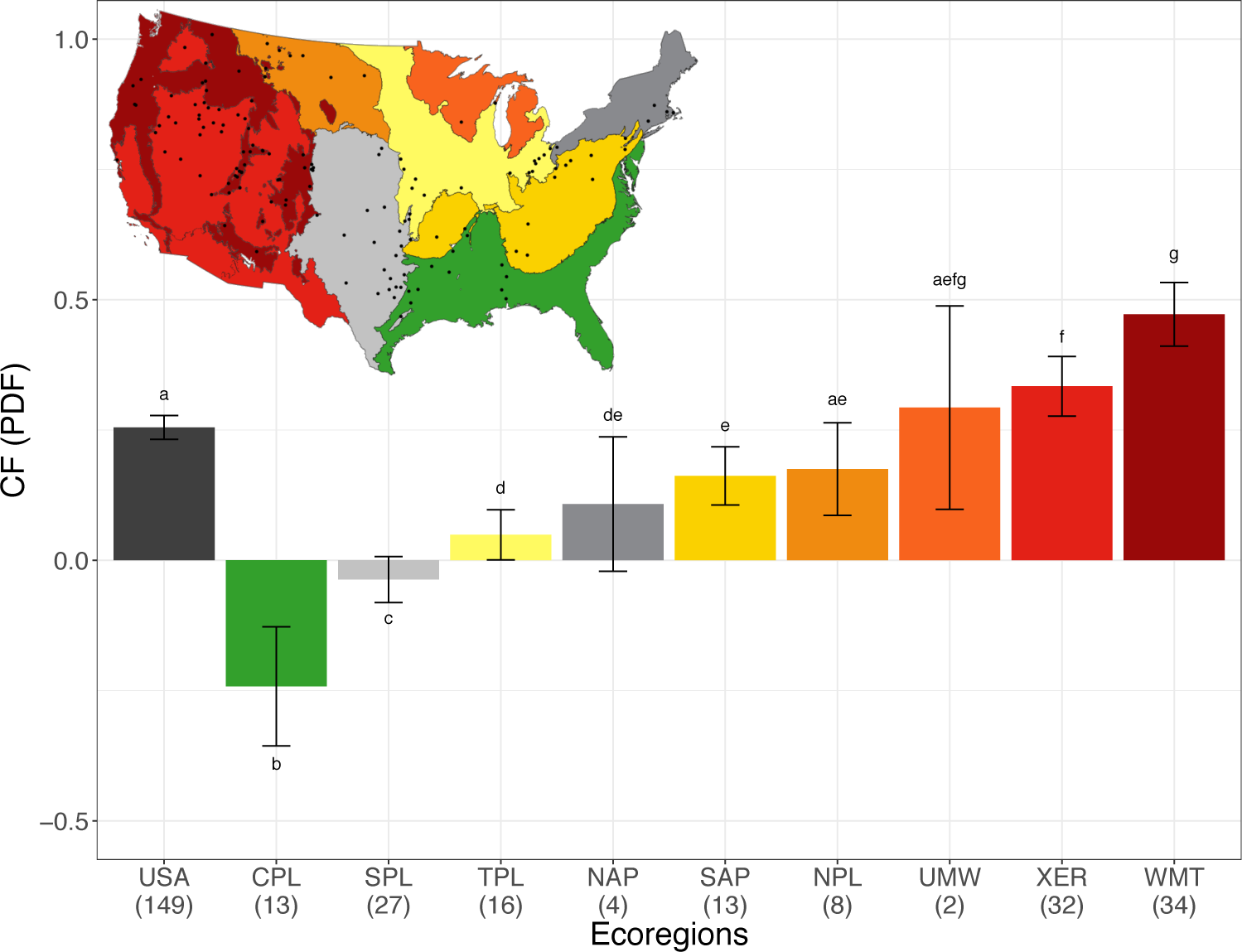
Barplot showing a mean PDF (± 95% confidence intervals) at the USA level (shown in dark grey) and at the ecoregion level (color-coded with ecoregions as a gradient of intensity). We used letter to identify which ecoregion mean PDF differ or not from each other. When two bars share the same letter, they are not significantly different from each other.

**Figure 3.**
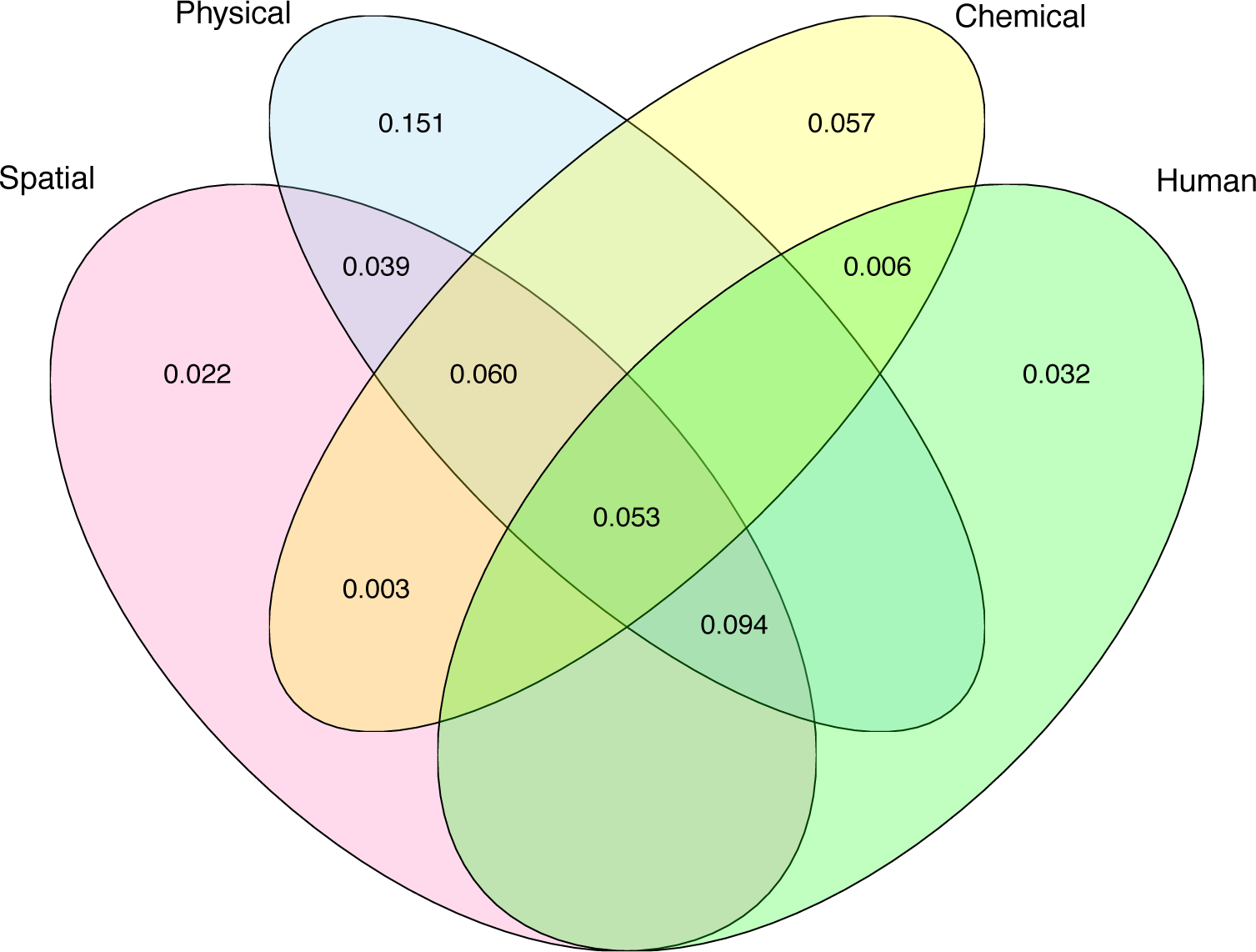
Venn diagram of the variation partitioning of a response matrix (PDF) explained by four matrices, that is spatial matrix (ECO), physical matrix (ELE + AREA), chemical matrix (T.S + PH) and human matrix (LAWN + ROAD). Values < 0 not shown.

**Table 2.** Table showing the mean PDF (± 95% confidence intervals) values for the USA and the nine ecoregions. Sample number is also shown (*n*).

| Ecoregion | PDF.m<sup>2</sup>.yr/m<sup>2</sup>.yr | ± 95% CIs | $n$ |
| --- | --- | --- | --- |
| USA | 0.255 | 0.123 | 148 |
| CPL | -0.242 | 0.182 | 13 |
| SPL | -0.037 | 0.101 | 27 |
| TPL | 0.049 | 0.085 | 16 |
| NAP | 0.108 | 0.114 | 4 |
| SAP | 0.162 | 0.089 | 13 |
| NPL | 0.175 | 0.111 | 8 |
| UMW | 0.293 | 0.122 | 2 |
| XER | 0.334 | 0.143 | 32 |
| WMT | 0.472 | 0.157 | 34 |

### 3.2 Variables explaining the variation in PDFs

At the reservoir scale, the four matrices (*i.e.*, spatial, physical, chemical, and human) explained 50% of the total variation observed in PDFs (variation partitioning; Figure 3; Table A2 for details on the percentages). About 42% of the variation was explained by the combined effects of the spatial (*i.e.*, ecoregion) and physical (*i.e.*, area of the reservoir and its elevation) matrices. The ecoregion (in the spatial matrix) explained 27% of the variation, over which 25% of this variation was shared with the physical matrix, 12% was shared with the chemical matrix (*i.e.*, pH and trophic state) and 14% was shared with the human matrix (*i.e.*, presence of lawn and road adjacent to the reservoir shoreline). The physical matrix explained 40% of the variation. Elevation and reservoir area alone (variation not shared with the other matrices) explained 15% of the variation. The chemical and human matrices explained respectively 18% and 19% of the variation.

### 3.3 PDFs predictive model

According to the predictive models (Equations 4 and 5; Figure 4) 50% of the observed variation in PDFs (partial R^2^_adj_ = 0.434; *p*-value < 0.001) was explained by elevation (33%), trophic state (*i.e.*, oligotrophic or eutrophic reservoir; 11%), and reservoir surface area (< 1%). PDF was positively related to reservoir elevation; the higher the elevation, the higher the PDF (Figure 4). PDF was also negatively related to eutrophication status. PDFs were higher in oligotrophic reservoirs (<10μg/l total phosphorus), and lower in eutrophic reservoirs (>10μg/l total phosphorus). As for reservoir surface area, there was a positive relationship between reservoir surface area and PDF; bigger reservoir had a higher PDF than smaller ones (Figure 4). In other words, large oligotrophic reservoirs located at higher elevation were most likely to have higher macroinvertebrate PDF.

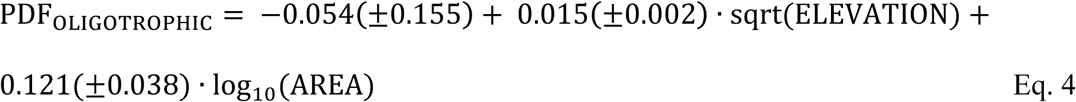

Values in parentheses are standard error (SE) of the estimate.

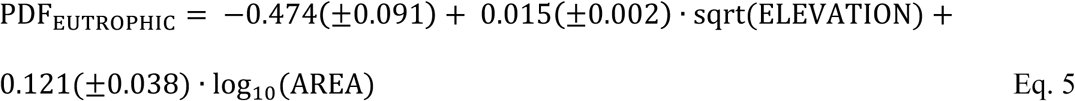

**Figure 4.**
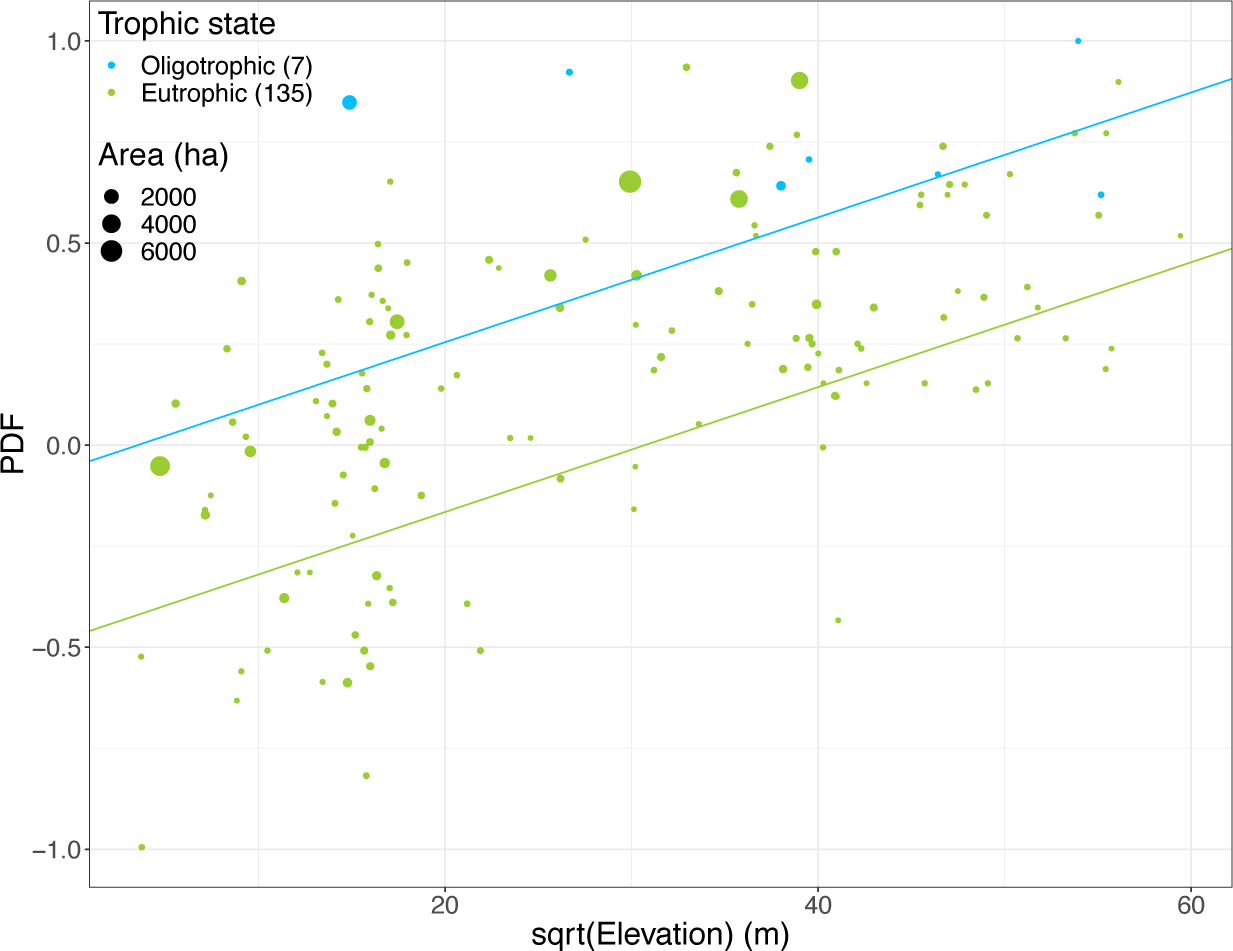
Graphical representation of our best empirical model showing the relationship between potentially disappeared fraction of species (PDF.m^2^.yr/m^2^.yr) and reservoir elevation (square-rooted). Trophic state is color-coded and point size is representative of reservoir surface area.

## 4. DISCUSSION

From the analysis of 148 reservoirs and 2121 rivers and streams across the continental United States, our results showed a loss of about 26% of macroinvertebrate taxa following impoundment at the scale of the United States. This result is consistent with the literature estimates of the impacts of hydropower on macroinvertebrate richness across the world (Kraft 1988, Englund and Malmqvist 1996, Malmqvist and Englund 1996, Jackson et al. 2007, Valdovinos et al. 2007, Takao et al. 2008, Aroviita and Hämäläinen 2008, White et al. 2011, Behrend et al. 2012, Kullasoot et al. 2017). PDFs also varied across ecoregions. More than 25% of the total variation observed in PDFs was explained by the nine ecoregions, pressing the need for regionalized CFs. We provided evidence that the empirical PDFs values for macroinvertebrates were consistent across our three spatial scales (country, ecoregions and reservoirs). Overall, our empirical PDFs can be used as CFs by LCA practitioners to evaluate the potential impact of reservoir occupation on the ecosystem quality area of protection. We also provided a simple predictive model based on three explanatory variables (*i.e.*, elevation, trophic state and reservoir surface area), and explaining 50% of the variation in macroinvertebrate PDFs. Reservoirs at higher elevation, with lower levels of eutrophication and bigger surface area were most likely to have higher PDFs. This predictive model could be used by LCA modelers to extrapolate CFs based on few explicative variables.

### 4.1 United States taxa loss and regionalization

At the scale of the United States, 26% of macroinvertebrate taxa disappeared following impoundment. The observed loss of native riverine macroinvertebrate taxa is an expected consequence of transforming a river into a reservoir. Macroinvertebrate taxa are highly adapted to their local habitat conditions (McCafferty 1983, Wetzel 2001). If flow regime is altered, which is a key characteristic of local riverine habitat, we can expect a substantial loss of these river specific taxa (Bunn & Arthington, 2002; Agostinho et al., 2008). The observed loss of taxa also followed a longitudinal gradient. PDFs are higher in the western than in the eastern ecoregions. The higher loss of taxa in the reservoirs of the “Western Mountains” (WMT) and “Xeric” (XER) ecoregions might be due to the higher water stress conditions. Reference rivers in these ecoregions are characterized by a higher proportion of intermittent streams, especially in dryer environment (Levick et al. 2008, USEPA 2013). This intermittent nature leads to highly variable environmental conditions, which increases taxonomic and functional biodiversity (Stubbington et al. 2017) and can lead to a potentially higher loss of taxa in reservoirs. Another complementary explanation to this higher PDF in the west could be related to reservoir topography and elevation (Figure A1). Reservoirs created in low relief environment can connect more waterbodies and watercourse than higher relief environment and can result in smaller PDFs, or even gains in taxa following impoundment (Gido et al. 2002, Gubiani et al. 2010). This enhanced water body connectivity can lead to a higher colonization pool, a higher resilience of the community, and can result in a lower loss of taxa following reservoir creation. This is very likely the case in the CPL ecoregion, which appears to be dominated by flat plains and a low relief topography (USEPA 2016), and is also the only ecoregion showing a potential appearance of taxa following reservoir creation. This ecoregion is also characterized by a high percentage of wetlands (USEPA, 2016), which can also enhance water body connectivity, and lead to negative PDF (*i.e.*, gain of taxa). Because information regarding these variables (*i.e.*, water stress/intermittence, waterbody connectivity, general topography and wetland presence) was not readily available from the datasets, we cannot statistically validate their relevance. Nonetheless, these significant differences in ecoregions’ PDFs strengthen the need for regionalization and the examination of multiple scales when developing CFs.

### 4.2 Trophic state

While elevation explained most of the variation in PDF, oligotrophic reservoirs (trophic state) had higher PDFs. One plausible explanation for this difference can be related to the low productivity nature of oligotrophic water bodies (Wetzel 2001). Considering that oligotrophic ecosystems sustain relatively low diversity (Dodson et al. 2000), oligotrophic reservoir PDFs are expected to be higher than eutrophic ones when compared with a mean ecoregion reference. The seven oligotrophic reservoirs vary in elevation (221 m to over 3000 m) and in surface area (9 ha to over 1900 ha). Eutrophic reservoirs showed a similar range in elevation and surface area, respectively 13 m to 3105 m and 4 ha to 6559 ha. This difference in PDF between oligotrophic and eutrophic reservoirs seems independent of these two variables. However, this observation was based on only seven oligotrophic reservoirs, versus 135 eutrophic reservoirs, a rather small and unbalanced sample size. We also observed that all oligotrophic reservoirs were located in the western part of the United States, the majority (*i.e.*, six out of seven) in the WMT (4) and XER (2) ecoregions. This observation could then be the result of a statistical artifact regarding the importance of ecoregion and could impose a limitation on the robustness of our model. Thus, caution is in order when it comes to interpreting this result.

### 4.3 Surface area

Reservoir surface area was also identified as an influential variable explaining variation in PDFs. While most of the reservoir surface areas ranged between 0 to 1500 ha, six particularly big reservoirs, between 2000 and 6000 ha, show higher loss of macroinvertebrates. Even though this trend is significant, it is quite weak considering the unbalanced sample size of big reservoirs versus average-sized reservoirs and the small variation explained (1%). These big reservoirs were also not clustered in one particular region of the United States or ecoregion, but well distributed in north-south and west-east gradients (Figure 1). While the majority of averaged sized reservoir were located between zero to over 3000 m of elevation, the six big reservoirs were located between zero and 1500m. They did not share a similar water usage (*e.g.*, hydropower, irrigation, recreation, etc.) and thus, no hypotheses can be drawn from the potential water extraction dynamic (*i.e.*, timing, rate, magnitude and frequency) or reservoirs usage on biodiversity impacts. One last plausible explanation could be related to reservoir bathymetry. Big shallow reservoirs usually have a substantial littoral zone (Zohary et al. 2014), which typically sustains higher macroinvertebrate productivity and diversity than its pelagic counterpart (James et al. 1998, Wetzel 2001). Water levels management (*e.g.*, fluctuations) in big shallow reservoirs can especially affect the survival of the macroinvertebrate community, thus resulting in higher loss of taxa (Aroviita and Hämäläinen 2008, White et al. 2011, Zohary et al. 2014). No bathymetry data was available from the datasets.

### 4.4 Limitations

In this study, we defined richness as the number of native riverine taxa. We did not account for the potential gain of lentic specific taxa following impoundment. When a river is transformed into a reservoir, some riverine species are lost, and some lentic taxa are gained. This gain in species, as well as gain in exotic and non-native invasive taxa should not be considered as an ecosystem improvement. Based on the habitat diversity hypothesis (Williams 1964), we can advocate that lotic environments are more diversified than lake-like ones (*i.e.*, reservoirs). This hypothesis states that the diversity of species is directly related to the diversity of habitats. Thus, because of their narrowness and longitude, rivers run through a greater range of geological formations as well as geographical regions per unit of surface area, and vary more in terms of substrate, water temperature and flow dynamics than lotic environment of comparable depth and size (Horwitz 1978, Moyle and Li 1979, Eadie et al. 1986). The increased environmental variability and productivity, as well as the presence of micro-habitat heterogeneity in rivers is likely to support more species per surface area (Gorman and Karr 1978, Matthews 1982, Eadie et al. 1986). We could then think that even after a lotic environment is transformed into and a lentic one, there would still be less taxa in the lentic environment. Moreover, gain of lentic species after impoundment is often considered a misleading argument because the littoral zone in reservoirs is less complex, differs in physico-chemical conditions (Walker et al. 1992) and is generally affected by varying water levels. Those characteristics can affect the productivity of littoral areas, which are crucial to reservoir productivity, and can, in turn, affect its biodiversity. Another limitation of this study is that our characterization model is not independent from other impact categories, namely eutrophication. Because trophic state was defined as a significant variable to predict PDF, we need to incorporate this information in our model. In the LCA framework, eutrophication is already taken into account and thus, using it in our model could cause some bias in the overall compilation of impacts (*i.e.*, double-counting). Finally, one last limitation from this study is the use of space-for-time substitution. We do not have a real before-after control impact study design. It was impossible to have access to river richness before impoundment so our results and suggested PDFs must be interpreted with caution.

## 5. CONCLUSION

In this study, by using space-for-time substitution, we showed that the transformation of a river into a reservoir resulted in a loss of 26% of macroinvertebrate taxa in the continental United States. This loss of richness also varied across ecoregions, pressing the need for regionalized CFs. Patterns were also consistent across scales (the United States, nine ecoregions and 148 reservoirs) where we observed a general loss of macroinvertebrate richness. These PDFs are destined to be used by LCA practitioners as CFs. We also derived a predictive model to estimate PDFs as a function of three explanatory variables; reservoir elevation, trophic state (*i.e.*, oligotrophic or eutrophic) and surface area. Despite a few highlighted limitations, the empirical CFs developed through this study constitute a strong contribution to assess the impacts of river impoundment on the ecosystem quality area of protection. A natural follow-up to this study would be to integrate macroinvertebrate CFs with fish CFs, from Turgeon et al. (2019, *submitted*; available as a preprint) to build more robust CFs. Moreover, we also recommend evaluating the environmental relevance of the predictive models in other geographical contexts. These steps would be advances toward a more unified and global coverage of the impacts of reservoir occupation on its ecosystem biodiversity.

## ACKNOWLEDGEMENTS

This work was financially supported by the Natural Sciences and Engineering Research Council of Canada (NSERCC), the Fonds Québécois de la Recherche sur la Nature et les technologies (FQRNT), Fondation Polytechnique and Hydro-Québec, as well as the Institut de l’Environnement, le Développement Durable et l’Économie Circulaire (EDDEC) and Banque TD. Funding sources had no involvement in the study design, data collection, analysis and interpretation, writing process and the decisions regarding manuscript submission for eventual journal publication.

## 7. APPENDICES

**Table A1.** Raw data.

| UID | LAT | LON | TROPHIC.STATE | ECO | N.REF | MEAN.REF.S | SD.REF.S | N.IMP | IMP.S | SD.IMP.S | DELTA.S | SD.POOLED | PDF | SD.PDF | LOW.CI | UP.CI |
| --- | --- | --- | --- | --- | --- | --- | --- | --- | --- | --- | --- | --- | --- | --- | --- | --- |
| 6243 | 38.50796545 | -94.67326537 | EUTROPHIC | TPL | 16 | 30.2 | 14.5 | 1 | 42 | 0 | -11.77 | 3.62 | -0.39 | -0.22 | -0.61 | -0.17 |
| 6252 | 40.1117666 | -75.8615303 | EUTROPHIC | SAP | 13 | 43.8 | 17.5 | 1 | 35 | 0 | 8.77 | 4.86 | 0.2 | 0.14 | 0.06 | 0.34 |
| 6267 | 31.62154629 | -88.35373896 | EUTROPHIC | CPL | 13 | 27.6 | 17.1 | 1 | 32 | 0 | -4.43 | 4.74 | -0.16 | -0.2 | -0.36 | 0.04 |
| 6270 | 35.17438787 | -99.07748881 | EUTROPHIC | SPL | 27 | 25.9 | 13.3 | 1 | 14 | 0 | 11.85 | 2.57 | 0.46 | 0.26 | 0.2 | 0.72 |
| 6281 | 35.28501346 | -112.1541634 | EUTROPHIC | WMT | 34 | 39.4 | 12.5 | 1 | 15 | 0 | 24.42 | 2.15 | 0.62 | 0.2 | 0.41 | 0.82 |
| 6319 | 36.83163302 | -104.2263032 | EUTROPHIC | WMT | 34 | 39.4 | 12.5 | 1 | 16 | 0 | 23.42 | 2.15 | 0.59 | 0.2 | 0.4 | 0.79 |
| 6342 | 39.99496234 | -105.112227 | EUTROPHIC | SPL | 27 | 25.9 | 13.3 | 1 | 26 | 0 | -0.15 | 2.57 | -0.01 | -0.1 | -0.11 | 0.09 |
| 6404 | 39.63835365 | -95.45681141 | EUTROPHIC | TPL | 16 | 30.2 | 14.5 | 1 | 22 | 0 | 8.23 | 3.62 | 0.27 | 0.18 | 0.1 | 0.45 |
| 6437 | 39.0001536 | -95.77964833 | EUTROPHIC | TPL | 16 | 30.2 | 14.5 | 1 | 34 | 0 | -3.77 | 3.62 | -0.12 | -0.13 | -0.26 | 0.01 |
| 6451 | 40.16178738 | -79.05238419 | EUTROPHIC | SAP | 13 | 43.8 | 17.5 | 1 | 43 | 0 | 0.77 | 4.86 | 0.02 | 0.11 | -0.09 | 0.13 |
| 6481 | 39.48435525 | -118.7235713 | EUTROPHIC | XER | 32 | 30.7 | 11.9 | 1 | 19 | 0 | 11.69 | 2.1 | 0.38 | 0.16 | 0.22 | 0.54 |
| 6482 | 41.92898525 | -119.1790015 | EUTROPHIC | XER | 32 | 30.7 | 11.9 | 1 | 16 | 0 | 14.69 | 2.1 | 0.48 | 0.2 | 0.28 | 0.68 |
| 6501 | 35.98082795 | -108.9316434 | EUTROPHIC | WMT | 34 | 39.4 | 12.5 | 1 | 14 | 0 | 25.42 | 2.15 | 0.64 | 0.21 | 0.43 | 0.86 |
| 6525 | 35.99293158 | -96.87367696 | EUTROPHIC | SPL | 27 | 25.9 | 13.3 | 1 | 40 | 0 | -14.15 | 2.57 | -0.55 | -0.3 | -0.85 | -0.25 |
| 6550 | 34.95361555 | -96.71815923 | EUTROPHIC | SPL | 27 | 25.9 | 13.3 | 1 | 27 | 0 | -1.15 | 2.57 | -0.04 | -0.1 | -0.15 | 0.06 |
| 6556 | 35.41220255 | -95.92927589 | EUTROPHIC | SPL | 27 | 25.9 | 13.3 | 1 | 25 | 0 | 0.85 | 2.57 | 0.03 | 0.1 | -0.07 | 0.13 |
| 6570 | 37.65556473 | -98.26098551 | EUTROPHIC | SPL | 27 | 25.9 | 13.3 | 1 | 39 | 0 | -13.15 | 2.57 | -0.51 | -0.28 | -0.79 | -0.23 |
| 6575 | 40.72381895 | -109.183908 | EUTROPHIC | XER | 32 | 30.7 | 11.9 | 1 | 21 | 0 | 9.69 | 2.1 | 0.32 | 0.14 | 0.18 | 0.46 |
| 6586 | 31.7872742 | -96.06449217 | EUTROPHIC | CPL | 13 | 27.6 | 17.1 | 1 | 28 | 0 | -0.43 | 4.74 | -0.02 | -0.17 | -0.19 | 0.16 |
| 6599 | 46.54392165 | -104.028258 | EUTROPHIC | NPL | 8 | 28.5 | 11.2 | 1 | 33 | 0 | -4.52 | 3.97 | -0.16 | -0.15 | -0.31 | -0.01 |
| 6606 | 37.39088653 | -99.78483802 | EUTROPHIC | SPL | 27 | 25.9 | 13.3 | 1 | 28 | 0 | -2.15 | 2.57 | -0.08 | -0.11 | -0.19 | 0.03 |
| 6617 | 41.75740963 | -115.7220267 | EUTROPHIC | XER | 32 | 30.7 | 11.9 | 1 | 26 | 0 | 4.69 | 2.1 | 0.15 | 0.09 | 0.06 | 0.24 |
| 6618 | 41.19805974 | -115.8922956 | EUTROPHIC | XER | 32 | 30.7 | 11.9 | 1 | 26 | 0 | 4.69 | 2.1 | 0.15 | 0.09 | 0.06 | 0.24 |
| 6622 | 31.88917209 | -97.70249202 | EUTROPHIC | SPL | 27 | 25.9 | 13.3 | 1 | 35 | 0 | -9.15 | 2.57 | -0.35 | -0.21 | -0.56 | -0.15 |
| 6667 | 43.37681333 | -123.2640104 | NA | WMT | 34 | 39.4 | 12.5 | 1 | 30 | 0 | 9.42 | 2.15 | 0.24 | 0.09 | 0.15 | 0.33 |
| 6668 | 36.82300133 | -96.04758798 | EUTROPHIC | SPL | 27 | 25.9 | 13.3 | 1 | 38 | 0 | -12.15 | 2.57 | -0.47 | -0.26 | -0.73 | -0.21 |
| 6695 | 36.70544343 | -96.4191085 | EUTROPHIC | TPL | 16 | 30.2 | 14.5 | 1 | 40 | 0 | -9.77 | 3.62 | -0.32 | -0.2 | -0.52 | -0.13 |
| 6719 | 38.39816173 | -115.117053 | EUTROPHIC | XER | 32 | 30.7 | 11.9 | 1 | 23 | 0 | 7.69 | 2.1 | 0.25 | 0.12 | 0.13 | 0.37 |
| 6731 | 41.70168986 | -113.9596706 | EUTROPHIC | XER | 32 | 30.7 | 11.9 | 1 | 26 | 0 | 4.69 | 2.1 | 0.15 | 0.09 | 0.06 | 0.24 |
| 6735 | 32.94442993 | -96.45375244 | EUTROPHIC | SPL | 27 | 25.9 | 13.3 | 1 | 34 | 0 | -8.15 | 2.57 | -0.32 | -0.19 | -0.51 | -0.12 |
| 6742 | 47.76171554 | -108.4328294 | EUTROPHIC | NPL | 8 | 28.5 | 11.2 | 1 | 30 | 0 | -1.52 | 3.97 | -0.05 | -0.14 | -0.19 | 0.09 |
| 6753 | 39.93104164 | -104.9732957 | EUTROPHIC | SPL | 27 | 25.9 | 13.3 | 1 | 20 | 0 | 5.85 | 2.57 | 0.23 | 0.15 | 0.07 | 0.38 |
| 6762 | 33.51617531 | -94.12513187 | EUTROPHIC | CPL | 13 | 27.6 | 17.1 | 1 | 43 | 0 | -15.43 | 4.74 | -0.56 | -0.39 | -0.95 | -0.17 |
| 6774 | 46.82604195 | -100.6342083 | EUTROPHIC | NPL | 8 | 28.5 | 11.2 | 1 | 16 | 0 | 12.48 | 3.97 | 0.44 | 0.22 | 0.22 | 0.66 |
| 6795 | 38.99724122 | -108.0511795 | EUTROPHIC | WMT | 34 | 39.4 | 12.5 | 1 | 32 | 0 | 7.42 | 2.15 | 0.19 | 0.08 | 0.11 | 0.27 |
| 6796 | 37.19334598 | -95.98897607 | EUTROPHIC | SPL | 27 | 25.9 | 13.3 | 1 | 36 | 0 | -10.15 | 2.57 | -0.39 | -0.23 | -0.62 | -0.17 |
| 6806 | 38.49141175 | -79.31478115 | EUTROPHIC | SAP | 13 | 43.8 | 17.5 | 1 | 43 | 0 | 0.77 | 4.86 | 0.02 | 0.11 | -0.09 | 0.13 |
| 6823 | 43.16587842 | -115.6524756 | EUTROPHIC | XER | 32 | 30.7 | 11.9 | 1 | 24 | 0 | 6.69 | 2.1 | 0.22 | 0.11 | 0.11 | 0.33 |
| 6825 | 37.25249786 | -113.7781188 | EUTROPHIC | XER | 32 | 30.7 | 11.9 | 1 | 2 | 0 | 28.69 | 2.1 | 0.93 | 0.37 | 0.57 | 1.3 |
| 6868 | 38.23508748 | -112.4630091 | EUTROPHIC | WMT | 34 | 39.4 | 12.5 | 1 | 26 | 0 | 13.42 | 2.15 | 0.34 | 0.12 | 0.22 | 0.46 |
| 6869 | 38.8475374 | -111.9613903 | EUTROPHIC | XER | 32 | 30.7 | 11.9 | 1 | 16 | 0 | 14.69 | 2.1 | 0.48 | 0.2 | 0.28 | 0.68 |
| 6874 | 39.03670301 | -107.9111308 | EUTROPHIC | WMT | 34 | 39.4 | 12.5 | 1 | 30 | 0 | 9.42 | 2.15 | 0.24 | 0.09 | 0.15 | 0.33 |
| 6875 | 40.94491896 | -106.0119684 | EUTROPHIC | XER | 32 | 30.7 | 11.9 | 1 | 26 | 0 | 4.69 | 2.1 | 0.15 | 0.09 | 0.06 | 0.24 |
| 6923 | 40.03999071 | -81.01388786 | EUTROPHIC | SAP | 13 | 43.8 | 17.5 | 1 | 24 | 0 | 19.77 | 4.86 | 0.45 | 0.21 | 0.24 | 0.66 |
| 6940 | 44.32909632 | -116.1841073 | EUTROPHIC | WMT | 34 | 39.4 | 12.5 | 1 | 29 | 0 | 10.42 | 2.15 | 0.26 | 0.1 | 0.16 | 0.36 |
| 6944 | 39.1695073 | -111.4507206 | EUTROPHIC | WMT | 34 | 39.4 | 12.5 | 1 | 29 | 0 | 10.42 | 2.15 | 0.26 | 0.1 | 0.16 | 0.36 |
| 6959 | 44.96411515 | -116.4630194 | EUTROPHIC | WMT | 34 | 39.4 | 12.5 | 1 | 32 | 0 | 7.42 | 2.15 | 0.19 | 0.08 | 0.11 | 0.27 |
| 6966 | 39.14241052 | -111.452546 | EUTROPHIC | WMT | 34 | 39.4 | 12.5 | 1 | 9 | 0 | 30.42 | 2.15 | 0.77 | 0.25 | 0.52 | 1.02 |
| 6970 | 44.79670477 | -116.7326881 | OLIGOTROPHIC | WMT | 34 | 39.4 | 12.5 | 1 | 13 | 0 | 26.42 | 2.15 | 0.67 | 0.22 | 0.45 | 0.89 |
| 6971 | 43.1914128 | -116.9598035 | EUTROPHIC | XER | 32 | 30.7 | 11.9 | 1 | 8 | 0 | 22.69 | 2.1 | 0.74 | 0.29 | 0.44 | 1.03 |
| 6976 | 38.7911491 | -105.1063606 | EUTROPHIC | WMT | 34 | 39.4 | 12.5 | 1 | 4 | 0 | 35.42 | 2.15 | 0.9 | 0.29 | 0.61 | 1.19 |
| 7001 | 42.2158744 | -83.58440358 | EUTROPHIC | TPL | 16 | 30.2 | 14.5 | 1 | 48 | 0 | -17.77 | 3.62 | -0.59 | -0.31 | -0.89 | -0.28 |
| 7020 | 38.07832575 | -122.7433592 | EUTROPHIC | XER | 32 | 30.7 | 11.9 | 1 | 36 | 0 | -5.31 | 2.1 | -0.17 | -0.1 | -0.27 | -0.08 |
| 7057 | 39.20473742 | -111.6689123 | EUTROPHIC | WMT | 34 | 39.4 | 12.5 | 1 | 30 | 0 | 9.42 | 2.15 | 0.24 | 0.09 | 0.15 | 0.33 |
| 7097 | 38.78818748 | -111.7748779 | EUTROPHIC | WMT | 34 | 39.4 | 12.5 | 1 | 15 | 0 | 24.42 | 2.15 | 0.62 | 0.2 | 0.41 | 0.82 |
| 7100 | 43.96521803 | -122.6839683 | EUTROPHIC | WMT | 34 | 39.4 | 12.5 | 1 | 37 | 0 | 2.42 | 2.15 | 0.06 | 0.06 | 0 | 0.12 |
| 7105 | 41.11051556 | -82.08387214 | EUTROPHIC | NAP | 4 | 44.6 | 15.7 | 1 | 28 | 0 | 16.58 | 7.87 | 0.37 | 0.22 | 0.15 | 0.59 |
| 7108 | 30.9634384 | -95.90350405 | EUTROPHIC | CPL | 13 | 27.6 | 17.1 | 1 | 27 | 0 | 0.57 | 4.74 | 0.02 | 0.17 | -0.15 | 0.19 |
| 7109 | 32.0726955 | -97.12977293 | EUTROPHIC | SPL | 27 | 25.9 | 13.3 | 1 | 24 | 0 | 1.85 | 2.57 | 0.07 | 0.11 | -0.03 | 0.18 |
| 7136 | 41.6332907 | -118.3893572 | EUTROPHIC | XER | 32 | 30.7 | 11.9 | 1 | 23 | 0 | 7.69 | 2.1 | 0.25 | 0.12 | 0.13 | 0.37 |
| 7168 | 48.27818524 | -116.8311247 | NA | WMT | 34 | 39.4 | 12.5 | 1 | 35 | 0 | 4.42 | 2.15 | 0.11 | 0.07 | 0.05 | 0.18 |
| 7205 | 32.24025447 | -101.3133027 | OLIGOTROPHIC | SPL | 27 | 25.9 | 13.3 | 1 | 2 | 0 | 23.85 | 2.57 | 0.92 | 0.49 | 0.44 | 1.41 |
| 7207 | 42.15782464 | -122.6076343 | EUTROPHIC | WMT | 34 | 39.4 | 12.5 | 1 | 26 | 0 | 13.42 | 2.15 | 0.34 | 0.12 | 0.22 | 0.46 |
| 7226 | 42.13001338 | -122.4782769 | EUTROPHIC | WMT | 34 | 39.4 | 12.5 | 1 | 19 | 0 | 20.42 | 2.15 | 0.52 | 0.17 | 0.34 | 0.69 |
| 7228 | 34.22781586 | -86.84344912 | EUTROPHIC | SAP | 13 | 43.8 | 17.5 | 1 | 44 | 0 | -0.23 | 4.86 | -0.01 | -0.11 | -0.12 | 0.11 |
| 7229 | 40.63158462 | -120.00287 | EUTROPHIC | XER | 32 | 30.7 | 11.9 | 1 | 20 | 0 | 10.69 | 2.1 | 0.35 | 0.15 | 0.2 | 0.5 |
| 7232 | 40.70340664 | -83.37874497 | EUTROPHIC | TPL | 16 | 30.2 | 14.5 | 1 | 17 | 0 | 13.23 | 3.62 | 0.44 | 0.24 | 0.2 | 0.68 |
| 7276 | 41.16858314 | -119.8174513 | OLIGOTROPHIC | XER | 32 | 30.7 | 11.9 | 1 | 9 | 0 | 21.69 | 2.1 | 0.71 | 0.28 | 0.42 | 0.99 |
| 7294 | 39.05604224 | -82.69067345 | EUTROPHIC | SAP | 13 | 43.8 | 17.5 | 1 | 47 | 0 | -3.23 | 4.86 | -0.07 | -0.12 | -0.19 | 0.04 |
| 7304 | 31.58749658 | -98.62250327 | EUTROPHIC | SPL | 27 | 25.9 | 13.3 | 1 | 36 | 0 | -10.15 | 2.57 | -0.39 | -0.23 | -0.62 | -0.17 |
| 7306 | 39.24114932 | -117.1658183 | EUTROPHIC | XER | 32 | 30.7 | 11.9 | 1 | 19 | 0 | 11.69 | 2.1 | 0.38 | 0.16 | 0.22 | 0.54 |
| 7325 | 40.33707955 | -105.1266943 | EUTROPHIC | SPL | 27 | 25.9 | 13.3 | 1 | 19 | 0 | 6.85 | 2.57 | 0.27 | 0.17 | 0.1 | 0.43 |
| 7367 | 40.58885713 | -111.5899784 | OLIGOTROPHIC | WMT | 34 | 39.4 | 12.5 | 1 | 0 | 0 | 39.42 | 2.15 | 1 | 0.32 | 0.68 | 1.32 |
| 7368 | 32.51586852 | -87.86108477 | EUTROPHIC | CPL | 13 | 27.6 | 17.1 | 1 | 29 | 0 | -1.43 | 4.74 | -0.05 | -0.17 | -0.23 | 0.12 |
| 7369 | 41.03542015 | -96.83772744 | EUTROPHIC | TPL | 16 | 30.2 | 14.5 | 1 | 26 | 0 | 4.23 | 3.62 | 0.14 | 0.14 | 0 | 0.28 |
| 7375 | 33.36486204 | -88.16688028 | EUTROPHIC | CPL | 13 | 27.6 | 17.1 | 1 | 45 | 0 | -17.43 | 4.74 | -0.63 | -0.43 | -1.06 | -0.2 |
| 7392 | 34.53435079 | -92.26882632 | EUTROPHIC | CPL | 13 | 27.6 | 17.1 | 1 | 26 | 0 | 1.57 | 4.74 | 0.06 | 0.18 | -0.12 | 0.23 |
| 7394 | 31.62154629 | -88.35373896 | EUTROPHIC | CPL | 13 | 27.6 | 17.1 | 1 | 28 | 0 | -0.43 | 4.74 | -0.02 | -0.17 | -0.19 | 0.16 |
| 7402 | 34.28477821 | -97.17097192 | EUTROPHIC | SPL | 27 | 25.9 | 13.3 | 1 | 39 | 0 | -13.15 | 2.57 | -0.51 | -0.28 | -0.79 | -0.23 |
| 7405 | 40.32809914 | -96.53200107 | EUTROPHIC | TPL | 16 | 30.2 | 14.5 | 1 | 25 | 0 | 5.23 | 3.62 | 0.17 | 0.15 | 0.03 | 0.32 |
| 7409 | 33.07556301 | -92.66059625 | EUTROPHIC | CPL | 13 | 27.6 | 17.1 | 1 | 31 | 0 | -3.43 | 4.74 | -0.12 | -0.19 | -0.31 | 0.06 |
| 7459 | 33.88200988 | -85.9316179 | EUTROPHIC | SAP | 13 | 43.8 | 17.5 | 1 | 39 | 0 | 4.77 | 4.86 | 0.11 | 0.12 | -0.01 | 0.23 |
| 7471 | 46.04062349 | -110.6921749 | EUTROPHIC | NPL | 8 | 28.5 | 11.2 | 1 | 23 | 0 | 5.48 | 3.97 | 0.19 | 0.16 | 0.03 | 0.35 |
| 7472 | 46.6246241 | -110.7383363 | EUTROPHIC | NPL | 8 | 28.5 | 11.2 | 1 | 25 | 0 | 3.48 | 3.97 | 0.12 | 0.15 | -0.03 | 0.27 |
| 7533 | 43.07839868 | -112.6936589 | EUTROPHIC | XER | 32 | 30.7 | 11.9 | 1 | 14 | 0 | 16.69 | 2.1 | 0.54 | 0.22 | 0.32 | 0.77 |
| 7572 | 40.37248248 | -84.34011043 | EUTROPHIC | TPL | 16 | 30.2 | 14.5 | 1 | 22 | 0 | 8.23 | 3.62 | 0.27 | 0.18 | 0.1 | 0.45 |
| 7579 | 39.60815568 | -84.97150743 | EUTROPHIC | TPL | 16 | 30.2 | 14.5 | 1 | 21 | 0 | 9.23 | 3.62 | 0.31 | 0.19 | 0.12 | 0.49 |
| 7587 | 42.26037727 | -111.4707171 | NA | WMT | 34 | 39.4 | 12.5 | 1 | 19 | 0 | 20.42 | 2.15 | 0.52 | 0.17 | 0.34 | 0.69 |
| 7588 | 42.89525373 | -111.9638056 | NA | XER | 32 | 30.7 | 11.9 | 1 | 36 | 0 | -5.31 | 2.1 | -0.17 | -0.1 | -0.27 | -0.08 |
| 7590 | 44.21474968 | -111.6155136 | EUTROPHIC | XER | 32 | 30.7 | 11.9 | 1 | 44 | 0 | -13.31 | 2.1 | -0.43 | -0.18 | -0.62 | -0.25 |
| 7596 | 40.86163552 | -75.61191223 | NA | SAP | 13 | 43.8 | 17.5 | 1 | 46 | 0 | -2.23 | 4.86 | -0.05 | -0.11 | -0.16 | 0.06 |
| 7643 | 39.70682362 | -111.2933692 | EUTROPHIC | WMT | 34 | 39.4 | 12.5 | 1 | 29 | 0 | 10.42 | 2.15 | 0.26 | 0.1 | 0.16 | 0.36 |
| 7652 | 40.17663921 | -84.26522022 | EUTROPHIC | TPL | 16 | 30.2 | 14.5 | 1 | 29 | 0 | 1.23 | 3.62 | 0.04 | 0.12 | -0.08 | 0.16 |
| 7684 | 37.67328141 | -107.1127777 | EUTROPHIC | WMT | 34 | 39.4 | 12.5 | 1 | 19 | 0 | 20.42 | 2.15 | 0.52 | 0.17 | 0.34 | 0.69 |
| 7686 | 37.31623199 | -107.1129943 | EUTROPHIC | WMT | 34 | 39.4 | 12.5 | 1 | 34 | 0 | 5.42 | 2.15 | 0.14 | 0.07 | 0.07 | 0.21 |
| 7698 | 41.15233931 | -110.8249531 | EUTROPHIC | XER | 32 | 30.7 | 11.9 | 1 | 8 | 0 | 22.69 | 2.1 | 0.74 | 0.29 | 0.44 | 1.03 |
| 7713 | 46.11821631 | -113.37464 | EUTROPHIC | WMT | 34 | 39.4 | 12.5 | 1 | 26 | 0 | 13.42 | 2.15 | 0.34 | 0.12 | 0.22 | 0.46 |
| 7769 | 30.97184148 | -88.0527428 | EUTROPHIC | CPL | 13 | 27.6 | 17.1 | 1 | 55 | 0 | -27.43 | 4.74 | -0.99 | -0.64 | -1.63 | -0.35 |
| 7800 | 41.67757315 | -73.14469782 | EUTROPHIC | NAP | 4 | 44.6 | 15.7 | 1 | 51 | 0 | -6.42 | 7.87 | -0.14 | -0.18 | -0.33 | 0.04 |
| 7810 | 43.41399799 | -119.4104723 | EUTROPHIC | XER | 32 | 30.7 | 11.9 | 1 | 10 | 0 | 20.69 | 2.1 | 0.67 | 0.27 | 0.4 | 0.94 |
| 7812 | 35.5624593 | -93.63756822 | EUTROPHIC | SAP | 13 | 43.8 | 17.5 | 1 | 28 | 0 | 15.77 | 4.86 | 0.36 | 0.18 | 0.18 | 0.54 |
| 7994 | 32.66237175 | -97.61238491 | EUTROPHIC | SPL | 27 | 25.9 | 13.3 | 1 | 47 | 0 | -21.15 | 2.57 | -0.82 | -0.43 | -1.25 | -0.38 |
| 8016 | 33.8293043 | -109.090421 | EUTROPHIC | WMT | 34 | 39.4 | 12.5 | 1 | 17 | 0 | 22.42 | 2.15 | 0.57 | 0.19 | 0.38 | 0.76 |
| 8121 | 36.06720319 | -91.14242843 | EUTROPHIC | SAP | 13 | 43.8 | 17.5 | 1 | 26 | 0 | 17.77 | 4.86 | 0.41 | 0.2 | 0.21 | 0.6 |
| 8144 | 35.58318947 | -90.96294056 | EUTROPHIC | CPL | 13 | 27.6 | 17.1 | 1 | 21 | 0 | 6.57 | 4.74 | 0.24 | 0.23 | 0.01 | 0.46 |
| 8151 | 41.08846075 | -82.7290152 | EUTROPHIC | TPL | 16 | 30.2 | 14.5 | 1 | 26 | 0 | 4.23 | 3.62 | 0.14 | 0.14 | 0 | 0.28 |
| 8184 | 40.05558648 | -105.7470802 | EUTROPHIC | WMT | 34 | 39.4 | 12.5 | 1 | 17 | 0 | 22.42 | 2.15 | 0.57 | 0.19 | 0.38 | 0.76 |
| 8207 | 39.65347461 | -82.47378134 | EUTROPHIC | SAP | 13 | 43.8 | 17.5 | 1 | 44 | 0 | -0.23 | 4.86 | -0.01 | -0.11 | -0.12 | 0.11 |
| 8250 | 48.38062147 | -110.9852659 | EUTROPHIC | NPL | 8 | 28.5 | 11.2 | 1 | 20 | 0 | 8.48 | 3.97 | 0.3 | 0.18 | 0.12 | 0.48 |
| 8256 | 39.77597099 | -81.52247236 | EUTROPHIC | SAP | 13 | 43.8 | 17.5 | 1 | 36 | 0 | 7.77 | 4.86 | 0.18 | 0.13 | 0.05 | 0.31 |
| 8278 | 48.02656863 | -109.6237599 | EUTROPHIC | NPL | 8 | 28.5 | 11.2 | 1 | 27 | 0 | 1.48 | 3.97 | 0.05 | 0.14 | -0.09 | 0.19 |
| 8325 | 37.4167824 | -108.4056506 | EUTROPHIC | WMT | 34 | 39.4 | 12.5 | 1 | 14 | 0 | 25.42 | 2.15 | 0.64 | 0.21 | 0.43 | 0.86 |
| 8342 | 32.05630851 | -96.73178299 | EUTROPHIC | SPL | 27 | 25.9 | 13.3 | 1 | 34 | 0 | -8.15 | 2.57 | -0.32 | -0.19 | -0.51 | -0.12 |
| 8360 | 40.67460225 | -110.9706993 | OLIGOTROPHIC | WMT | 34 | 39.4 | 12.5 | 1 | 15 | 0 | 24.42 | 2.15 | 0.62 | 0.2 | 0.41 | 0.82 |
| 8395 | 40.88926752 | -109.8461084 | EUTROPHIC | WMT | 34 | 39.4 | 12.5 | 1 | 24 | 0 | 15.42 | 2.15 | 0.39 | 0.14 | 0.26 | 0.53 |
| 8409 | 39.72076713 | -86.72022287 | EUTROPHIC | TPL | 16 | 30.2 | 14.5 | 1 | 30 | 0 | 0.23 | 3.62 | 0.01 | 0.12 | -0.11 | 0.13 |
| 8413 | 31.91034371 | -95.30185594 | EUTROPHIC | CPL | 13 | 27.6 | 17.1 | 1 | 38 | 0 | -10.43 | 4.74 | -0.38 | -0.29 | -0.67 | -0.09 |
| 8414 | 40.12169493 | -104.9459108 | EUTROPHIC | SPL | 27 | 25.9 | 13.3 | 1 | 6 | 0 | 19.85 | 2.57 | 0.77 | 0.41 | 0.36 | 1.18 |
| 8416 | 38.93934838 | -91.28285684 | EUTROPHIC | TPL | 16 | 30.2 | 14.5 | 1 | 37 | 0 | -6.77 | 3.62 | -0.22 | -0.16 | -0.38 | -0.06 |
| 8427 | 44.69847871 | -87.49992572 | EUTROPHIC | UMW | 2 | 38.9 | 13.3 | 1 | 30 | 0 | 8.87 | 9.44 | 0.23 | 0.26 | -0.03 | 0.48 |
| 8435 | 47.87668819 | -107.1252315 | EUTROPHIC | NPL | 8 | 28.5 | 11.2 | 1 | 14 | 0 | 14.48 | 3.97 | 0.51 | 0.24 | 0.26 | 0.75 |
| 8437 | 42.64690672 | -72.21849668 | EUTROPHIC | NAP | 4 | 44.6 | 15.7 | 1 | 40 | 0 | 4.58 | 7.87 | 0.1 | 0.18 | -0.08 | 0.28 |
| 8443 | 41.81248656 | -70.63822658 | EUTROPHIC | CPL | 13 | 27.6 | 17.1 | 1 | 42 | 0 | -14.43 | 4.74 | -0.52 | -0.37 | -0.89 | -0.16 |
| 8480 | 30.00697302 | -96.70980987 | EUTROPHIC | SPL | 27 | 25.9 | 13.3 | 1 | 39 | 0 | -13.15 | 2.57 | -0.51 | -0.28 | -0.79 | -0.23 |
| 8487 | 39.66169251 | -84.64620693 | EUTROPHIC | TPL | 16 | 30.2 | 14.5 | 1 | 20 | 0 | 10.23 | 3.62 | 0.34 | 0.2 | 0.14 | 0.54 |
| 8494 | 41.98955896 | -71.20506636 | EUTROPHIC | NAP | 4 | 44.6 | 15.7 | 1 | 40 | 0 | 4.58 | 7.87 | 0.1 | 0.18 | -0.08 | 0.28 |
| 8495 | 46.94126809 | -119.27853 | EUTROPHIC | XER | 32 | 30.7 | 11.9 | 1 | 34 | 0 | -3.31 | 2.1 | -0.11 | -0.08 | -0.19 | -0.03 |
| 8504 | 38.07061898 | -111.3751268 | EUTROPHIC | WMT | 34 | 39.4 | 12.5 | 1 | 9 | 0 | 30.42 | 2.15 | 0.77 | 0.25 | 0.52 | 1.02 |
| 8614 | 36.04444348 | -85.58629482 | EUTROPHIC | SAP | 13 | 43.8 | 17.5 | 1 | 22 | 0 | 21.77 | 4.86 | 0.5 | 0.23 | 0.27 | 0.73 |
| 8627 | 48.687922 | -121.6679362 | OLIGOTROPHIC | WMT | 34 | 39.4 | 12.5 | 1 | 6 | 0 | 33.42 | 2.15 | 0.85 | 0.28 | 0.57 | 1.12 |
| 8635 | 40.49553571 | -83.89988027 | EUTROPHIC | TPL | 16 | 30.2 | 14.5 | 1 | 21 | 0 | 9.23 | 3.62 | 0.31 | 0.19 | 0.12 | 0.49 |
| 8765 | 37.59556184 | -112.2546686 | EUTROPHIC | WMT | 34 | 39.4 | 12.5 | 1 | 25 | 0 | 14.42 | 2.15 | 0.37 | 0.13 | 0.24 | 0.49 |
| 8766 | 40.86378075 | -109.8118146 | EUTROPHIC | WMT | 34 | 39.4 | 12.5 | 1 | 13 | 0 | 26.42 | 2.15 | 0.67 | 0.22 | 0.45 | 0.89 |
| 8777 | 39.36035119 | -111.9637079 | EUTROPHIC | XER | 32 | 30.7 | 11.9 | 1 | 3 | 0 | 27.69 | 2.1 | 0.9 | 0.36 | 0.55 | 1.26 |
| 8811 | 31.33182559 | -97.26843785 | EUTROPHIC | SPL | 27 | 25.9 | 13.3 | 1 | 41 | 0 | -15.15 | 2.57 | -0.59 | -0.32 | -0.9 | -0.27 |
| 1000019 | 43.46019017 | -116.1413012 | EUTROPHIC | XER | 32 | 30.7 | 11.9 | 1 | 25 | 0 | 5.69 | 2.1 | 0.19 | 0.1 | 0.09 | 0.28 |
| 1000025 | 43.19840666 | -114.5997261 | EUTROPHIC | XER | 32 | 30.7 | 11.9 | 1 | 27 | 0 | 3.69 | 2.1 | 0.12 | 0.08 | 0.04 | 0.2 |
| 1000029 | 42.67704536 | -113.4072962 | EUTROPHIC | XER | 32 | 30.7 | 11.9 | 1 | 12 | 0 | 18.69 | 2.1 | 0.61 | 0.25 | 0.36 | 0.85 |
| 1000030 | 42.20637168 | -114.8788521 | EUTROPHIC | XER | 32 | 30.7 | 11.9 | 1 | 20 | 0 | 10.69 | 2.1 | 0.35 | 0.15 | 0.2 | 0.5 |
| 1000062 | 33.27090609 | -98.15099199 | EUTROPHIC | SPL | 27 | 25.9 | 13.3 | 1 | 9 | 0 | 16.85 | 2.57 | 0.65 | 0.35 | 0.3 | 1 |
| 1000068 | 46.20607125 | -116.8343671 | EUTROPHIC | XER | 32 | 30.7 | 11.9 | 1 | 22 | 0 | 8.69 | 2.1 | 0.28 | 0.13 | 0.15 | 0.41 |
| 1000073 | 35.58232959 | -101.7171387 | EUTROPHIC | SPL | 27 | 25.9 | 13.3 | 1 | 9 | 0 | 16.85 | 2.57 | 0.65 | 0.35 | 0.3 | 1 |
| 1000084 | 41.03740677 | -100.7757749 | EUTROPHIC | SPL | 27 | 25.9 | 13.3 | 1 | 15 | 0 | 10.85 | 2.57 | 0.42 | 0.24 | 0.18 | 0.66 |
| 1000086 | 41.32198742 | -98.90067497 | EUTROPHIC | SPL | 27 | 25.9 | 13.3 | 1 | 15 | 0 | 10.85 | 2.57 | 0.42 | 0.24 | 0.18 | 0.66 |
| 1000088 | 41.76432182 | -98.65979155 | NA | SPL | 27 | 25.9 | 13.3 | 1 | 27 | 0 | -1.15 | 2.57 | -0.04 | -0.1 | -0.15 | 0.06 |
| 1000122 | 42.26425757 | -116.3104956 | EUTROPHIC | XER | 32 | 30.7 | 11.9 | 1 | 25 | 0 | 5.69 | 2.1 | 0.19 | 0.1 | 0.09 | 0.28 |
| 1000126 | 42.53147462 | -116.3645133 | EUTROPHIC | XER | 32 | 30.7 | 11.9 | 1 | 23 | 0 | 7.69 | 2.1 | 0.25 | 0.12 | 0.13 | 0.37 |
| 1000137 | 42.1873061 | -113.92408 | OLIGOTROPHIC | XER | 32 | 30.7 | 11.9 | 1 | 11 | 0 | 19.69 | 2.1 | 0.64 | 0.26 | 0.38 | 0.9 |
| 1000223 | 43.53712279 | -90.95935275 | EUTROPHIC | UMW | 2 | 38.9 | 13.3 | 1 | 25 | 0 | 13.87 | 9.44 | 0.36 | 0.27 | 0.08 | 0.63 |

**Table A2.**
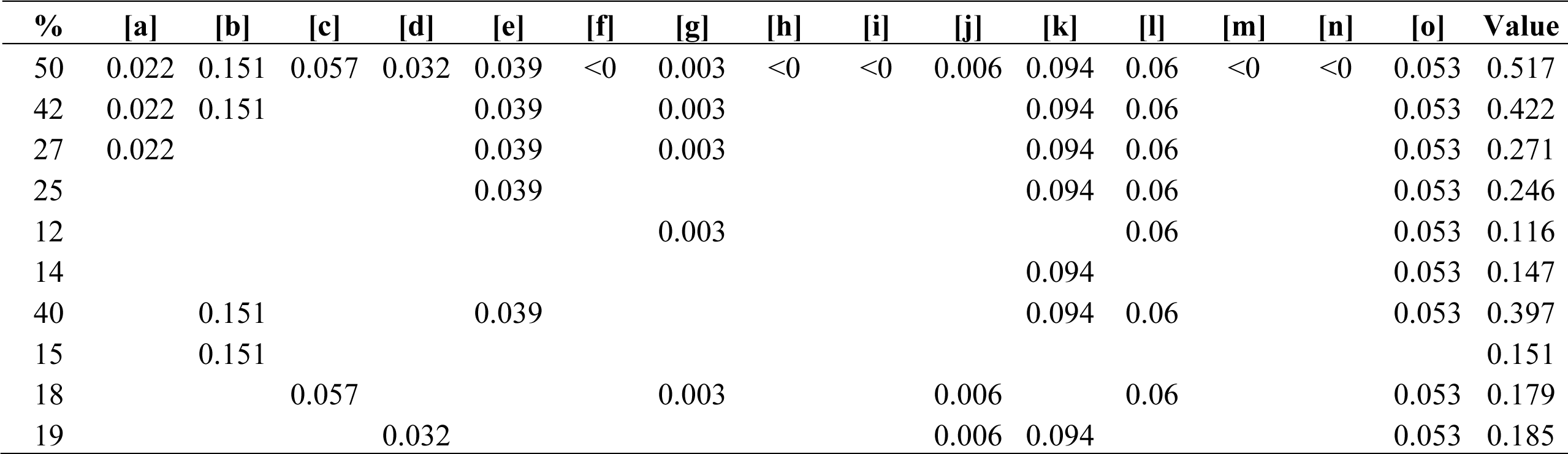
Variation partitioning fractions (*i.e.*, percentages) explained.

| % | [a] | [b] | [c] | [d] | [e] | [f] | [g] | [h] | [i] | [j] | [k] | [l] | [m] | [n] | [o] | Value |
| --- | --- | --- | --- | --- | --- | --- | --- | --- | --- | --- | --- | --- | --- | --- | --- | --- |
| 50 | 0.022 | 0.151 | 0.057 | 0.032 | 0.039 | <0 | 0.003 | <0 | <0 | 0.006 | 0.094 | 0.06 | <0 | <0 | 0.053 | 0.517 |
| 42 | 0.022 | 0.151 |  |  | 0.039 |  | 0.003 |  |  |  | 0.094 | 0.06 |  |  | 0.053 | 0.422 |
| 27 | 0.022 |  |  |  | 0.039 |  | 0.003 |  |  |  | 0.094 | 0.06 |  |  | 0.053 | 0.271 |
| 25 |  |  |  |  | 0.039 |  |  |  |  |  | 0.094 | 0.06 |  |  | 0.053 | 0.246 |
| 12 |  |  |  |  |  |  | 0.003 |  |  |  |  | 0.06 |  |  | 0.053 | 0.116 |
| 14 |  |  |  |  |  |  |  |  |  |  | 0.094 |  |  |  | 0.053 | 0.147 |
| 40 |  | 0.151 |  |  | 0.039 |  |  |  |  |  | 0.094 | 0.06 |  |  | 0.053 | 0.397 |
| 15 |  | 0.151 |  |  |  |  |  |  |  |  |  |  |  |  |  | 0.151 |
| 18 |  |  | 0.057 |  |  |  | 0.003 |  |  | 0.006 |  | 0.06 |  |  | 0.053 | 0.179 |
| 19 |  |  |  | 0.032 |  |  |  |  |  | 0.006 | 0.094 |  |  |  | 0.053 | 0.185 |

**Figure A1.**
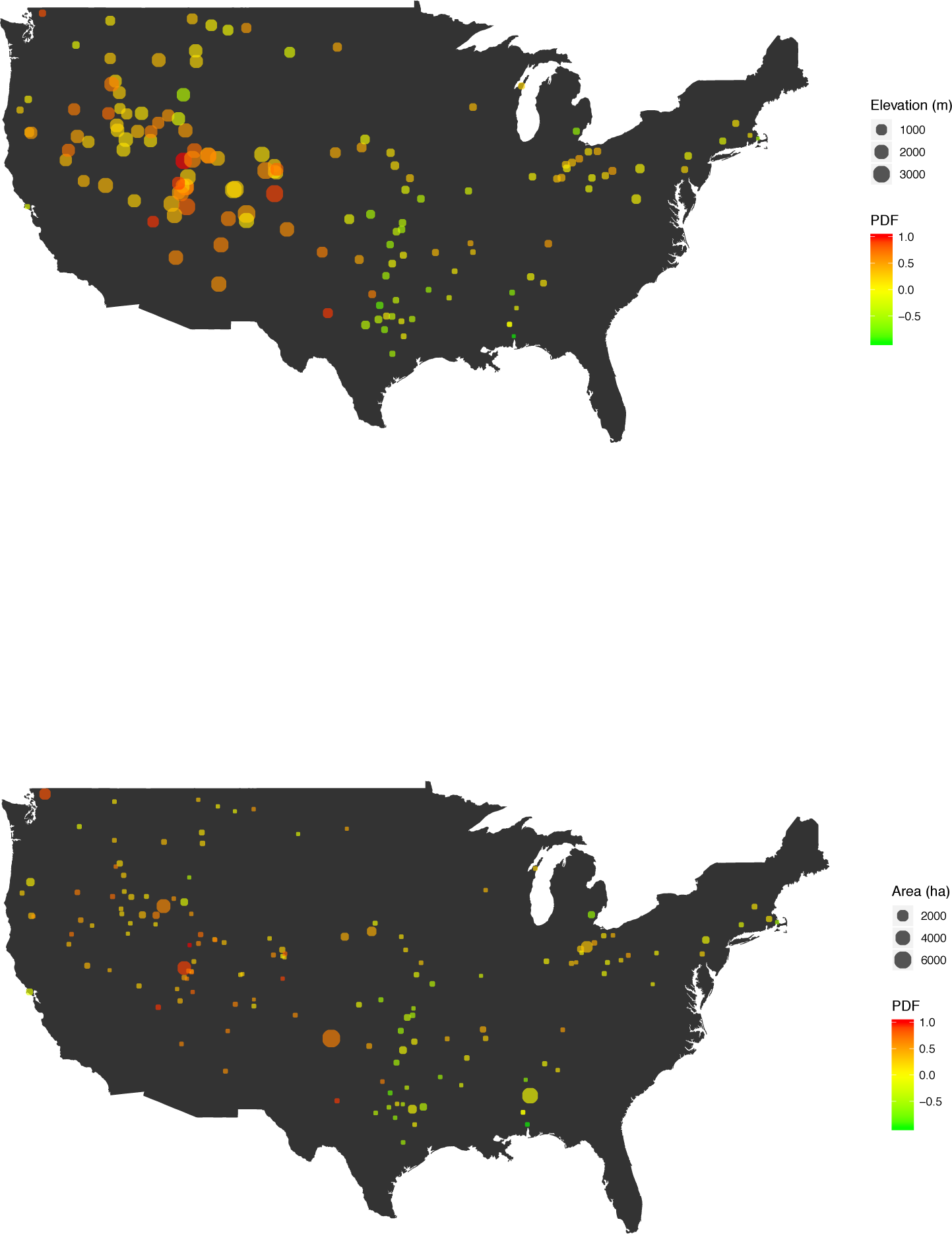
Heatmaps of PDF, elevation and surface area of reservoir is proportional to the point size.

